# Comparison of the performance of multiple whole-genome sequence-based tools for the identification of *Bacillus cereus sensu stricto* biovar Thuringiensis

**DOI:** 10.1101/2024.01.23.575246

**Authors:** Taejung Chung, Abimel Salazar, Grant Harm, Sophia Johler, Laura M. Carroll, Jasna Kovac

**Affiliations:** Department of Food Science, The Pennsylvania State University, University Park, PA; Institute for Food Safety and Hygiene, Vetsuisse Faculty, University of Zurich, Zurich, Switzerland; Department of Clinical Microbiology, SciLifeLab, Umeå University, Umeå, Sweden; Laboratory for Molecular Infection Medicine Sweden (MIMS), Umeå University, Umeå, Sweden; Umeå Centre for Microbial Research (UCMR), Umeå University, Umeå, Sweden; Integrated Science Lab (IceLab), Umeå University, Umeå, Sweden

**Keywords:** *Bacillus thuringiensis*, biopesticide, whole genome sequencing, Bt-toxin

## Abstract

The *Bacillus cereus sensu stricto* (*s.s.*) species comprises strains of biovar *Thuringiensis* (*Bt*) known for their bioinsecticidal activity, as well as strains with foodborne pathogenic potential. *Bt* strains are identified (i) based on the production of insecticidal crystal proteins also known as Bt toxins or (ii) based on the presence of *cry*, *cyt*, and *vip* genes, which encode Bt toxins. Multiple bioinformatics tools have been developed for the detection of crystal protein-encoding genes based on whole-genome sequencing (WGS) data. However, the performance of these tools is yet to be evaluated using phenotypic data. Thus, the goal of this study was to assess the performance of four bioinformatics tools for the detection of crystal protein-encoding genes. The accuracy of sequence-based identification of *Bt* was determined in reference to phenotypic microscope-based screening for production of crystal proteins. A total of 58 diverse *B. cereus s.l.* strains isolated from clinical, food, environmental, and commercial biopesticide products were underwent WGS. Isolates were examined for crystal protein production using phase contrast microscopy. Crystal protein-encoding genes were detected using BtToxin_Digger, BTyper3, IDOPS, and Cry_processor. Out of 58 isolates, the phenotypic production of crystal proteins was confirmed for 18 isolates. Specificity and sensitivity of *Bt* identification based on sequences were 0.85 and 0.94 for BtToxin_Digger, 0.97 and 0.89 for BTyper3, 0.95 and 0.94 for IDOPS, and 0.88 and 1.00 for Cry_processor, respectively. Cry_processor predicted crystal protein production with highest specificity, and BtToxin_Digger and IDOPS predicted crystal protein production with the highest sensitivity. Three out of four tested bioinformatic tools performed well overall, with IDOPS achieving both high sensitivity and specificity (>0.90).

**IMPORTANCE:** *Bacillus cereus s.s.* biovar *Thuringiensis* (*Bt*) is used as an organic biopesticide. It is differentiated from the foodborne pathogen *Bacillus cereus s.s.* by the production of insecticidal crystal proteins. Thus, reliable genomic identification of biovar *Thuringiensis* is necessary to ensure food safety and facilitate risk assessment. This study assessed the accuracy of WGS-based identification of *Bt* compared to phenotypic microscopy-based screening for crystal protein production. Multiple bioinformatics tools were compared to assess their performance in predicting crystal protein production. Among them, IDOPS performed best overall at WGS- based *Bt* identification.

## INTRODUCTION

The *Bacillus cereus* group, also known as *B. cereus sensu lato* (*s.l.*), is a species complex that comprises strains with the ability to cause human illness, as well as strains that have agriculturally and industrially beneficial phenotypes. Among well-known species in the *B. cereus* group are foodborne pathogen *B. cereus sensu stricto* (here referred to as *Bc*) (1) and entomopathogen *B. thuringiensis* (here referred to as *Bt*), the latter of which is commercially available as a biopesticide for application in organic farming (2).

The insecticidal activity of *Bt* is supported by the production of insecticidal proteins (i.e., Bt toxins), including parasporal crystal proteins such as crystal (Cry) and cytolytic (Cyt) toxins, and non-parasporal proteins such as vegetative insecticidal proteins (Vip) (3, 4). Bt toxins act against different insect species, including Lepidoptera, Diptera, Coleoptera, and Hymenoptera (3, 5). In 1995, the United States Environmental Protection Agency (US EPA) registered the first *Bt* biopesticide products for use in the US (6). Currently, over 180 *Bt* products are registered under 15 different EPA product code numbers, and they all include strains of at least one of the four *Bt* subspecies (i.e., *kurstaki*, *israelensis*, *aizawai*, and *tenebrionis*) (6). However, despite the long history of agricultural application of *Bt*, the concerns over safety of *Bt* for humans have been raised over the past decade (7–11).

*Bt* strains have been reported to encode and produce human enterotoxins that are known to contribute to foodborne illness caused by *Bc* (8, 9, 12). These include hemolysin BL (Hbl), non-hemolytic enterotoxin (Nhe), and cytotoxin K (CytK) (13–16). Given that routine diagnostic assays do not differentiate between *Bt* and *Bc*, it is possible that some *Bc-*associated outbreaks may have been caused by *Bt* (17). However, direct evidence for *Bt* causing human illness has not been explicitly established (17, 18).

The high genomic similarity of *Bc* and *Bt* results in their classification into the same genomospecies, even based on the least conservative classification criteria for genomospecies (13). Moreover, these two species cannot be distinguished by typical culture-based detection methods: the only phenotypic trait that is used for differentiating *Bc* and *Bt* is microscopy-based detection of parasporal crystal proteins (i.e. Bt toxins), which are responsible for the bioinsecticidal properties of *Bt* strains (19, 20) (21). The above-outlined taxonomic classification shortcomings have been addressed in a new proposed taxonomic framework that considers both genomic and phenotypic information for *B. cereus* group species identification (22). Within the proposed taxonomic framework, *Bc* and *Bt* are represented by one genomospecies named *Bacillus cereus sensu stricto*. Furthermore, a biovar *Thuringiensis* has been defined to represent genomes that carry crystal protein-encoding genes (i.e., Bt toxin genes), including *cry*, *cyt*, and *vip* genes (2). Reliable detection of Bt toxin genes is therefore required for accurate genome-based detection of *B. cereus s.l.* biovar *Thuringiensis* (*Bt*).

One of the main challenges in detecting Bt toxin genes is a high variability in their sequences. For example, over 300 variants of *cry* genes have been identified based on the amino acid sequence similarity (23, 24). Thus, multiple whole-genome sequencing (WGS)-based bioinformatics tools have been developed not only to detect known Bt toxin gene sequences, but also to predict new variants of these genes. These tools include BtToxin_Digger (25), IDOPS (26), Cry_processor (27), and BTyper3 (13). Despite the availability of these tools, their performance has not been compared by phenotypic assays. This study therefore aimed to (i) compare Bt toxin-encoding gene identification tools and (ii) compare their performance against phenotypic data (i.e., microscopy screening for crystal protein production).

## RESULTS AND DISCUSSION

We first compared the completeness of hybrid and short read assemblies using the N50 metric and the number of contigs, as determined by QUAST (**Figure 1**). N50 values for hybrid assemblies (min = 443,575 bp, max = 5,675,203 bp, mean = 4,245,832.3 bp) were significantly higher than N50 for short read assemblies (min = 22,040 bp, max = 514,228 bp, mean = 113,641.28 bp) (t-test p-value = 1.98 x 10^-47^). The number of contigs equal or longer than 1,000 bp was significantly higher in short read assemblies (min = 32, max = 607, mean = 235) compared to hybrid assemblies (min = 1, max = 98, mean = 17) (t-test p-value = 7.37 x 10^-23^). N50 values were, on average, 59% shorter in short-read assemblies than hybrid assemblies.

**Figure 1.**
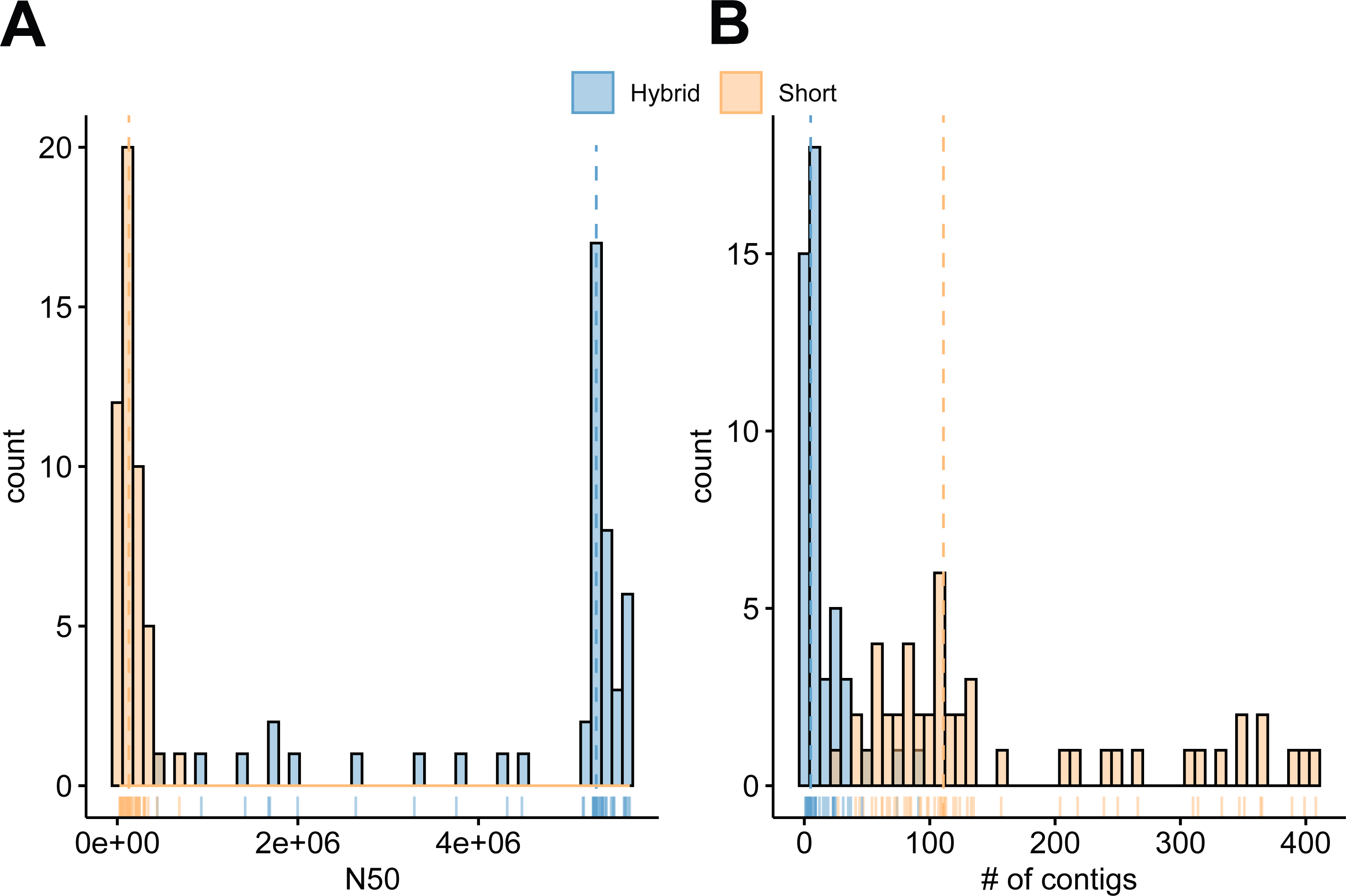
Distribution of contig counts in hybrid and short read assemblies. (A) N50 and (B) number of total contigs (=> 1000 bp).

Similarly, there were on average 79% less contigs in hybrid assemblies than short-read assemblies. Both of these metrics taken together indicate that hybrid assemblies were more complete. Both types of assemblies were further used for the assessment of bioinformatics tools for the detection of Bt toxin-encoding genes to measure the effect of assembly completeness on the performance of individual tools.

### IDOPS performed best overall for Bt toxin-encoding gene detection in comparison to phenotypic data

Prior to the application of bioinformatic tools for the detection of Bt toxin-encoding genes, we identified and removed redundant genomes (here, defined as genomes that differed by < 6 SNPs). Briefly, using PubMLST’s seven-gene multi-locus sequence typing (MLST) scheme for “*B. cereus*”, 37 different sequence types (STs) were identified, with biovar *Thuringiensis* identified in 10 STs (i.e., ST1085, ST1099, ST1142, ST138, ST15, ST1734, ST325, ST33, ST414, and ST8). Out of the 37 different MLST STs, 6 STs had more than one isolate (i.e., ST1099, ST1424, ST15, ST8, ST24, and ST73); thus, high-quality SNPs were identified within those STs. As a result, a total of 20 clonal genomes were excluded from further analyses to mitigate clonal redundancy bias. This resulted in a total of 58 isolates that were included in the assessment of the performance of four bioinformatics tools (i.e., IDOPS, Cry_processor, BtToxin_Digger, and BTyper3). IDOPS (Identification of Pesticidal Sequences) uses profile HMMs (Hidden Markov models) (28) to detect pesticidal protein-encoding genes in accordance with the BPPRC nomenclature system (Bacterial Pesticidal Protein Resource Center) (23). Cry_processor was developed based on HMM-based algorithm and has two different search modes available: domain only (DO) and find domain (FD). DO mode detects the sequences comprising all the three domains in the right order, and FD mode search novel domain of the matched Bt toxin gene sequences, using Bt-Toxin nomenclature database (29). Here, we tested both modes to compare the performance of the pipeline. There are no differences in results between two modes. BtToxin_Digger uses three different methods (i.e., BLAST, HMM, and support vector machine [SVM]) for the identification of Bt toxin gene sequences using the Bt toxin nomenclature database. Lastly, BTyper3 uses BLAST to detect Bt toxin genes using translated amino acid sequences with conservative detection thresholds of 70% identity and 50% coverage.

The production of crystal proteins was confirmed for 18 out of 58 isolates using phase contrast microscopy (**Figure 2**). When applied to short-read assemblies, CryProcessor predicted crystal protein production with highest specificity (0.97). BtToxin_Digger and IDOPS predicted crystal protein production with the highest sensitivity (0.94), whereas BtToxin_Digger had the lowest specificity (0.82). When applied to hybrid assemblies, the same bioinformatics programs showed highest and lowest specificities and sensitivities (**Table 1**). Specifically, CryProcessor predicted protein production with higher specificity (1.00) compared to the other three programs: BTyper3 (0.97), BtToxin_Digger (0.85), and IDOPS (0.95). However, the other three programs showed higher sensitivity than CryProcessor (0.83): BTyper3 (0.88), BtToxin _Digger (0.94), and IDOPS (0.94). Similar to results observed for short-read assemblies, BtToxin_Digger had the lowest specificity when applied to hybrid assemblies (0.82). These results indicate that genome assembly completeness was not a major factor influencing the performance of bioinformatics tools for the detection of Bt toxin-encoding genes. Three out of four tested bioinformatic tools (i.e., IDOPS, BTyper3, and Cry_processor) performed well overall, with IDOPS achieving sensitivity and specificity >0.90. BtToxin_Digger was highly sensitive but also non-specific, as it resulted in the highest number of false positive hits (*n* = 6). The other 3 tools produced a maximum of 3 false positive hits (**Table 1**).

**Figure 2.**
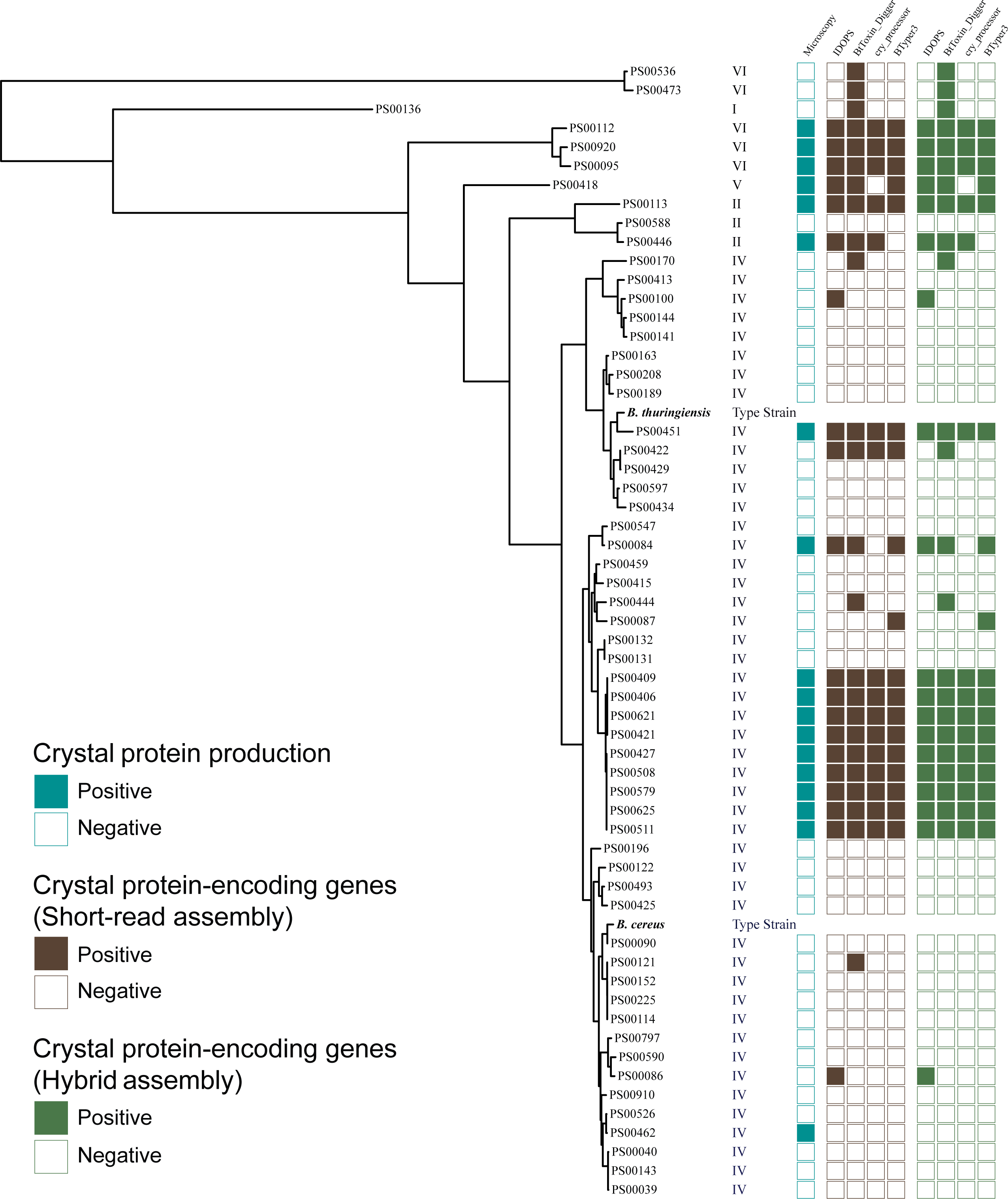
Maximum-likelihood phylogenetic tree for studied *B. cereus* group strains. Adjacent to the tips of the tree is a heatmap displaying prediction of crystal protein production based on the detected crystal-protein-encoding genes (*cry*, *cyt*, and *vip*), and a phylogenetic group for each isolate. The type strains of *B. cereus* and *B. Thuringiensis* were included as references.

**TABLE 1.**
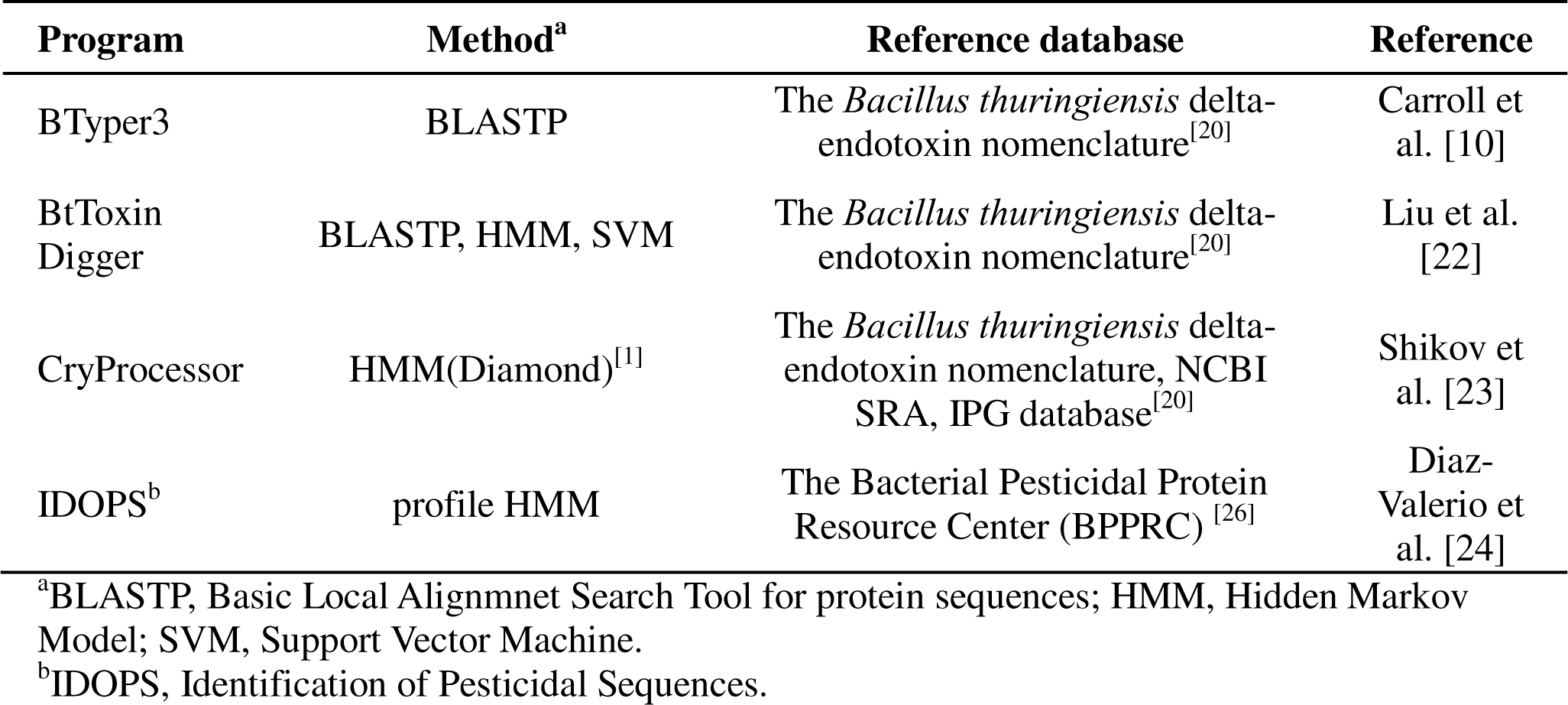
Description of bioinformatics tools for prediction of Bt-toxin-encoding genes.

Even though IDOPS performed best among the assessed bioinformatics tools, it made two false positives predictions (i.e., isolates PS00086 and PS00100). These two isolates were not phylogenetically closely related and were predicted as positive for Bt toxin-encoding genes by IDOPS’ profile HMMs 5 (CryM5) and 6 (CryM6). All other IDOPS models (i.e., CryM1 to M4, CytM1, CytM2, and VipM1) correctly predicted the presence of Bt toxin-encoding genes. The CryM5 profile HMM targets Cry proteins with less conserved variants of the classical 3 domains, while CryM6 targets the C-terminal region of Cry pesticidal proteins (26). This could explain why the CryM5 and CryM6 models produced false positive predictions on non-*Bt* strains.

BtToxin_Digger, which had the lowest specificity, uses a combination of multiple algorithms, including BLAST, HMM, and SVM. We examined whether false positive results were caused by any specific algorithm and found that the majority of falls positive calls were attributable to the outcomes of predictive models (i.e., HMM and SVM) (**Table 2**). HMM had both low sensitivity and specificity (Sn = 0.68, Sp = 0.68), whereas SVM had low sensitivity and high specificity (Sn = 0.00, Sp = 0.95). In contrast to HHM and SVM, we found that BLAST search alone performed very well (Sn = 0.97, Sp = 0.95) and produced BtToxin_Digger results that were comparable to the other three programs.

**TABLE 2.** Sensitivity, specificity, and positive and negative predictive values for predicting crystal protein production using four bioinformatics tools.

| **Assembly** | **Programs** | **Sensitivity** | **Specificity** | **PPV<sup>a</sup>** | **NPV<sup>b</sup>** |
| --- | --- | --- | --- | --- | --- |
| Hybrid | BTyper3 | 0.89 | 0.97 | 0.94 | 0.95 |
|  | BtToxin Digger | 0.94 | 0.85 | 0.73 | 0.97 |
|  | CryProcessor | 0.88 | 1.00 | 1 | 0.95 |
|  | IDOPS<sup>c</sup> | 0.94 | 0.95 | 0.89 | 0.97 |
| Short read | BTyper3 | 0.88 | 0.95 | 0.88 | 0.95 |
|  | BtToxin Digger | 0.94 | 0.82 | 0.70 | 0.97 |
|  | CryProcessor | 0.83 | 0.97 | 0.93 | 0.93 |
|  | IDOPS<sup>c</sup> | 0.94 | 0.92 | 0.85 | 0.97 |

<sup>a</sup>PPV, positive predictive value.
<sup>b</sup>NPV, negative predictive value.
<sup>c</sup>IDOPS, identification of pesticidal sequences.

### Phylogenetic distribution of biovar *Thuringiensis* strains within the *B. cereus* group

Overall, 12 out of 18 biovar *Thuringiensis* strains were classified into adjusted eight-group *panC* phylogenetic group IV, 2 into group II, 1 into group V, and 3 into group VI **(Figure 2)**. The maximum likelihood phylogenetic analysis showed that the biovar *Thuringiensis* strains (as defined by the detection of Bt toxin-encoding genes by at least 3 bioinformatics tools or the microscopic screening) were found across different lineages of phylogenetic group IV (corresponding to *B. cereus s.s.*) and were intermixed with non-*Thuringiensis* strains (**Figure 2**). Notably, most of the biovar *Thuringiensis* strains included in this study were not closely related to the *B. thuringinesis* type strain (ATCC 10792) nor the *B. cereus s.s.* type strain (ATCC 14579). Average SNP differences relative to the *B. thuringiensis* type strain were 44,765 SNPs (min = 7,717 SNPs, max = 76,229 SNPs); for the *B. cereus s.s.* type strain, strains differed by an average of 37,748.94 SNPs (min = 6,808 SNPs, max = 62,696 SNPs). This demonstrates that phylogenetic analysis alone is not sufficient for the identification of strains belonging to biovar *Thuringiensis*. This finding agrees with previous studies and suggests that Bt toxin genes (e.g., *cry, cyt,* and *vip*) likely disseminated among *B. cereus* group strains through horizontal gene transfer (16). Bt toxin genes are typically found on plasmids (30, 31), although they have also been detected within the chromosome of the *B. thuringiensis* HER1410 strain (32). We found Bt toxin-encoding genes on non-chromosomal contigs in 17 out of 18 hybrid assemblies of strains belonging to biovar *Thuringiensis*. Only one isolate (PS00095) carried the *cry27Aa1* gene on a chromosomal contig. This gene shared 75% identity and 100% coverage with the reference sequence in the *B. thuringiensis* delta-endotoxin nomenclature database. A transfer and chromosome integration of a plasmid-borne gene sequence may have been facilitated by a transposase; however, no transposable elements were detected adjacent to *cry27Aa1*.

## MATERIALS AND METHODS

### Isolates included in the study

A total of 78 *B. cereus* group strains available in the Kovac lab culture collection, some of which have been reported previously (15, 33), were included in the study based on the following criteria: (i) an isolate was classified as *B. cereus s.s.* (phylogenetic group IV) using BTyper 3 (v3.2.0) (n=72) or (ii) an isolate belonged to any of the *B. cereus s.l.* phylogenetic groups and had *cry, cyt,* and/or *vip* genes detected in its draft genome using BTyper3 (n=6) (v3.2.0) (13). Isolate information, including year of isolation, origin of isolation, and NCBI accession numbers, are listed in the Supplemental Material (**Table S1**).

### Illumina whole-genome sequencing and sequence quality control

Total genomic DNA was extracted from isolates using E.Z.N.A Bacterial DNA extraction kit (Omega Bio-Tek, GA, USA) by following the manufacturer’s instructions. Extracted DNA was examined for quality and quantity using Nanodrop One (Thermo Fisher Scientific, MA, USA) and Qubit 3 (Thermo Fisher Scientific, MA, USA), respectively. DNA libraries were prepared using a NexteraXT library preparation kit (Illumina, CA, USA). Samples were sequenced using an Illumina NextSeq with 150 bp paired-end reads. Additionally, genomes of 15 isolates with publicly available WGS reads were obtained from the NCBI Sequence Read Archive (SRA; NCBI BioProject Accessions PRJNA437714 or PRJNA288461) (15, 33). Illumina adapters and low-quality reads were trimmed using Trimmomatic (v0.39) with a sliding window size of 4 and quality cutoff value of 15 (SLIDINGWINDOW:4:15) (34). Reads shorter than 36 bp were excluded (MINLEN:36). Trimmed reads qualities were assessed using FastQC (v0.11.9) (35).

### Nanopore whole-genome sequencing and sequence quality control

DNA was extracted using the QIAamp DNA blood mini kit (Qiagen, Hilden, Germany) by following the manufacturer’s instructions with additional steps for cell lysis. Briefly, 3 loopfuls of biomass grown at 30°C on Brain-Heart Infusion (BHI) agar (BD Biosciences, NJ, USA) was collected and mixed with a phosphate-buffered saline (PBS) buffer (137 mM NaCl, 1.7 mM KCl, 10 mM Na_2_HPO_4_, 1.8 mM KH_2_PO_4_). One gram of 0.1 mm zirconia/silica beads (Biospec Products, OK, USA) were added to tubes, which were then vortexed in a horizontal position to disrupt cell walls. The Oxford Nanopore Technologies (ONT) RBK-004 rapid barcoding kit was used for the library preparation by following manufacturer’s instructions, and 4 libraries were pooled for sequencing (ONT, Oxford, UK). An R9 flow cell (FLO-MIN106; ONT, Oxford, UK) was used for sequencing using a MinION Mk1C device (ONT). Raw sequencing signal files were used for high accuracy basecalling and adapter trimming using Guppy (v6.0.1). Low-quality reads were trimmed using FiltLong (v0.2.1) (36), and the quality of trimmed reads was assessed using FastQC (v0.11.9).

### Genome assembly

Illumina reads were assembled *de novo* with SPAdes using the “isolate” mode (--isolate) and *k*-mer sizes of 99 or 127 (-k 99,127) (v3.15.3) (37). Trimmed Illumina short reads and trimmed ONT long reads were used for hybrid genome assembly with Unicycler using default parameters (v0.5.0) (38). Assembly quality was assessed using QUAST (v5.0.2) (39).

### Maximum-likelihood phylogenetic tree construction

To visualize phylogenetic relationships among isolates, core-genome single nucleotide polymorphisms (cgSNPs) of assembled genomes were identified by kSNP3 (v3.1.2) (40), using *k*=21; assembled genomes were used as input, along with the *B. cereus* ATCC 14579 and *B. thuringiensis* ATCC 10792 type strains genomes (GenBank Accession numbers GCA_018309165 and GCA_000161615, respectively). Maximum-likelihood (ML) phylogenetic trees were constructed using the detected cgSNPs and IQ-TREE 2 (v2.2.0), using the generalized time-reversible model with a gamma distribution and ascertainment bias correction (GTR+G+ASC), plus 1000 ultrafast bootstraps (41–44). The final ML tree was visualized and annotated using iTOL (v6.8) (45).

To remove redundant and clonal isolates, high-quality SNPs were identified using the FDA CFSAN SNP pipeline with default setting (v.2.2.1) (46). This analysis was conducted separately for isolates belonging to each individual MLST sequence type (ST) that contained two or more isolates. MLST STs were assigned using BTyper 3 (v3.2.0) and PubMLST’s seven-gene MLST scheme for “*B. cereus*” (47). Pairwise SNP differences were calculated within each ST, and for each group of clonal genomes (i.e., isolates that differed by < 6 whole-genome SNPs), one high-quality representative genome was selected based on N50 values and number of contigs. The same SNP pipeline was applied to all identified biovar *Thuringiensis* genomes by using two type strains as references (*B. thuringiensis* type strain ATCC 10792 and *B. cereus* type strain ATCC 14579) to calculate average SNP differences between biovar *Thuringiensis* strains and the two type strains.

### Bioinformatic detection of Bt-toxin-encoding genes

Bt toxin-encoding genes were detected in each assembled genome using each of the following bioinformatic pipelines: BTyper3 (v3.2.0) (13), BtToxin_Digger (v1.0.10) (25), IDOPS (v0.2.2) (26), and Cry_processor (Last update at August 2019; domain only [DO] and find domain [FD] were used) (27) (**Table 3**). The four bioinformatic tools were applied to both short-read and hybrid assemblies. Protein coding sequences (CDS) were identified using Prokka with default setting (v1.14.5) (48). The resulting CDSs were used as an input for IDOPS, Cry_processor, and BtToxin_Digger. Nucleotide genome assemblies were used as an input for BTyper3.

**TABLE 3.** Sensitivity, specificity, and positive and negative predictive values for predicting crystal protein production using different BtToxin_digger algorithms (i.e., BLAST, HMM model, SVM model) applied on hybrid assemblies.

| **Algorithm<sup>a</sup>** | **Sensitivity** | **Specificity** | **PPV<sup>b</sup>** | **NPV<sup>b</sup>** |
| --- | --- | --- | --- | --- |
| BLASTP | 0.97 | 0.95 | 0.94 | 0.98 |
| HMM | 0.68 | 0.68 | 0.62 | 0.73 |
| SVM | 0.00 | 0.95 | 0.00 | 0.55 |

<sup>a</sup>BLASTP, Basic Local Alignment Search Tool for protein sequences; HMM, Hidden Markov Model; SVM, Support Vector Machine.
<sup>b</sup>PPV, positive predictive value; NPV, negative predictive value.

### Microscopy-based screening of isolates for production of insecticidal crystal proteins

Isolates were grown on T3 agar for 3 days at 30°C to promote sporulation and crystal protein production. After completed incubation, one colony was resuspended in 10 µL of PBS on a microscope slide and covered with a cover slip (VWR international, PA, USA) and screened within 10 minutes for the presence of crystal proteins using phase-contrast microscope (Olympus BX51, Olympus, Tokyo, Japan). Crystal proteins were differentiated by shape (bipyramidal, cuboidal, and circular), size (smaller than a vegetative cell and endospore of *Bacillus*), and color (darker than *Bacillus* endospore). Initial screening results were classified as “positive”, “negative”, or “inconclusive”. Screening was repeated three times by two individuals to assess the production of crystal proteins.

### Statistical analysis

All statistical analyses in the study were conducted using R (version 4.2.1) (49). A paired t-test was employed to compare the average N50 and the number of contigs between short-read assembly and hybrid assembly (50). The sensitivity, specificity, positive prediction value (PPV), and negative prediction value (NPV) were calculated to assess the performance of four different bioinformatic tools for the detection of Bt toxin-encoding genes (51). Genome assemblies produced using short reads, as well as and hybrid assemblies were used. The results of microscopy screening were used to validate the sequence-based bioinformatic prediction of *B. cereus* biovar *Thuringiensis*.

## ACKNOWLEDGEMENTS

This work was supported by the USDA National Institute of Food and Agriculture and Hatch Appropriations under Project PEN04853 and Accession 7005519, project 4666, and USDA Agriculture and Food Research Initiative Research and Extension Experiences for Undergraduates grant 2021-67037-34628 “Engaging Students from Undergraduate-Centric Institutions in Research, Informatics, and Experiential Opportunities in Food Microbiology”. LMC was supported by the SciLifeLab & Wallenberg Data Driven Life Science Program (grant: KAW 2020.0239).

## DATA AVAILABILITY STATEMENT

Paired-end Illumina reads sequenced in this study have been deposited in the NCBI SRA database under BioProject accession number PRJNA715191. Single-end Nanopore sequences have been deposited in the NCBI SRA database under BioProject accession number PRJNA1007391. Detailed information for sequencing data used in this study is available in Table S1. Scripts for bioinformatic analyses are available in the GitHub repository (https://github.com/tuc289/Bt_toxin_tools_validation).

## Notes

### Competing Interest Statement

The authors have declared no competing interest.

