## Supplemental Table 1 for "Comparison of the performance of multiple whole-genome sequence-based tools for the identification of *Bacillus cereus sensu stricto* biovar Thuringiensis"

**Table S1.** Name, phylogenetic groups, sources, and NCBI SRA accessions for isolates used in this study.

| **Isolate name** | **Phylogenetic group** | **Isolate source** | **NCBI SRA accession for short read sequences** | **NCBI SRA accession for long read sequences** |
| --- | --- | --- | --- | --- |
| PS00039 | IV | food | SRR6827983 | SRR25793468 |
| PS00040 | IV | unknown | SRR6827982 | SRR25793467 |
| PS00084 | IV | food | SRR18750957 | SRR25793445 |
| PS00086 | IV | food | SRR4661783 | SRR25793434 |
| PS00087 | IV | food | SRR23629687 | SRR25793423 |
| PS00090 | IV | unknown | SRR5189060 | SRR25793412 |
| PS00095 | VI | food | SRR4661786 | SRR25813338 |
| PS00100 | IV | food | SRR23629676 | SRR25793401 |
| PS00112 | VI | environment | SRR23629671 | SRR25813337 |
| PS00113 | II | food | SRR18750928 | SRR25813334 |
| PS00114 | IV | food | SRR5189061 | SRR25793391 |
| PS00121 | IV | food | SRR5185012 | SRR25793390 |
| PS00122 | IV | food | SRR18750966 | SRR25793466 |
| PS00131 | IV | food | SRR18750962 | SRR25793465 |
| PS00132 | IV | food | SRR18750961 | SRR25793464 |
| PS00136 | I | environment | SRR4661789 | SRR25813333 |
| PS00141 | IV | environment | SRR23629667 | SRR25793463 |
| PS00143 | IV | food | SRR2541693 | SRR25793462 |
| PS00144 | IV | food | SRR5185011 | SRR25793461 |
| PS00152 | IV | food | SRR5185023 | SRR25793460 |
| PS00163 | IV | food | SRR2541604 | SRR25793459 |
| PS00170 | IV | food | SRR18750952 | SRR25793458 |
| PS00189 | IV | food | SRR5185020 | SRR25793455 |
| PS00196 | IV | food | SRR5185019 | SRR25793453 |
| PS00208 | IV | food | SRR23629694 | SRR25793451 |
| PS00225 | IV | food | [SRR21459607](https://dataview.ncbi.nlm.nih.gov/object/39382322) | SRR25813332 |
| PS00406 | IV | food | SRR13988996 | SRR25793443 |
| PS00409 | IV | food | SRR13988963 | SRR25793440 |
| PS00413 | IV | food | SRR13988908 | SRR25793436 |
| PS00415 | IV | food | SRR13988906 | SRR25793435 |
| PS00418 | V | food | SRR13988903 | SRR25813331 |
| PS00421 | IV | food | SRR13988896 | SRR25813330 |
| PS00422 | IV | food | SRR13988896 | SRR25793433 |
| PS00425 | IV | food | SRR13988892 | SRR25793431 |
| PS00427 | IV | food | SRR13988890 | SRR25793430 |
| PS00429 | IV | food | SRR13988888 | SRR25793428 |
| PS00434 | IV | food | SRR13988882 | SRR25793425 |
| PS00444 | IV | food | SRR13988871 | SRR25793422 |
| PS00446 | II | food | SRR13988869 | SRR25813329 |
| PS00451 | IV | food | SRR13988867 | SRR25793421 |
| PS00459 | IV | food | SRR13988861 | SRR25793420 |
| PS00462 | IV | food | SRR13988860 | SRR25793419 |
| PS00473 | VI | food | SRR13988851 | SRR25813328 |
| PS00493 | IV | environment | SRR13988833 | SRR25793418 |
| PS00508 | IV | biopesticide | SRR13989016 | SRR25793413 |
| PS00511 | IV | biopesticide | SRR13989012 | SRR25793409 |
| PS00526 | IV | food | SRR13988999 | SRR25793407 |
| PS00536 | VI | food | SRR13988990 | SRR25813327 |
| PS00547 | IV | food | SRR13988977 | SRR25793406 |
| PS00579 | IV | food | SRR13988949 | SRR25793402 |
| PS00588 | II | food | SRR23629675 | SRR25813336 |
| PS00590 | IV | food | SRR13988940 | SRR25793400 |
| PS00597 | IV | food | SRR13988937 | SRR25793399 |
| PS00621 | IV | food | SRR18750940 | SRR25793397 |
| PS00625 | IV | food | SRR13988885 | SRR25793396 |
| PS00797 | IV | food | SRR13988964 | SRR25793393 |
| PS00910 | IV | food | SRR13988922 | SRR25793392 |
| PS00920 | VI | food | SRR13988917 | SRR25813335 |
